# Temporal Dynamics of Normalization Reweighting

**DOI:** 10.1101/2023.02.10.527994

**Authors:** Daniel H. Baker, Daniela Marinova, Richard Aveyard, Lydia J. Hargreaves, Alice Renton, Ruby Castellani, Phoebe Hall, Miriam Harmens, Georgia Holroyd, Beth Nicholson, Emily L. Williams, Hannah M. Hobson, Alex R. Wade

## Abstract

For decades, neural suppression in early visual cortex has been thought to be fixed. But recent work has challenged this assumption by showing that suppression can be *reweighted* based on recent history; when pairs of stimuli are repeatedly presented together, suppression between them strengthens. Here we investigate the temporal dynamics of this process using a steady-state visual evoked potential (SSVEP) paradigm that provides a time-resolved, direct index of suppression between pairs of stimuli flickering at different frequencies (5 and 7Hz). Our initial analysis of an existing EEG dataset (N=100) indicated that suppression increases substantially during the first 2-5 seconds of stimulus presentation (with some variation across stimulation frequency). We then collected new EEG data (N=100) replicating this finding for both monocular and dichoptic mask arrangements in a preregistered study designed to measure reweighting. A third experiment (N=20) used source localized MEG, and found that these effects are apparent in primary visual cortex (V1), consistent with results from neurophysiological work. Because long-standing theories propose inhibition/excitation differences in autism, we also compared reweighting between individuals with high vs low autistic traits, and with and without an autism diagnosis, across our 3 data sets (total N=220). We find no compelling differences in reweighting that are associated with autism. Our results support the normalization reweighting model, and indicate that for prolonged stimulation, increases in suppression occur on the order of 2-5 seconds after stimulus onset.

## 2 Introduction

Suppressive interactions between neurons are ubiquitous in the nervous system, with normalization considered a canonical neuronal computation (Carandini & Heeger, 2011). One consequence of normalization is that neurons tuned to different stimulus features modulate each others’ firing, usually via a process of divisive suppression (Heeger, 1992). For decades the strength of suppression was treated as fixed, due to the observation that adapting to one stimulus does not decrease its suppressive potency (Foley & Chen, 1997; Freeman et al., 2002). This orthodoxy has been challenged by a series of innovative studies proposing that normalization can be ‘reweighted’ by recent history (Aschner et al., 2018; Westrick et al., 2016; Yiltiz et al., 2020). When pairs of stimuli are repeatedly presented together, their neural representations appear to suppress each other more strongly. Far from being fixed, normalization may therefore be a dynamic process that is continuously updated by the sensory environment. Our objectives were to measure the time-course of changes in suppression non-invasively in the human brain, compare them across distinct anatomical pathways, and determine whether they differ as a function of autistic traits.

Plastic changes within the visual system occur over multiple timescales (see Webster, 2015 for a recent review). Cortical forms of adaptation to cues such as stimulus contrast (Blakemore & Campbell, 1969), orientation (Gibson & Radner, 1937) and motion (Mather et al., 2008) can be observed within a few seconds, but also build up over durations on the order of minutes (Greenlee et al., 1991). Other types of adaptation have been identified where changes occur over longer time periods, such as several hours (Kwon et al., 2009) or days (Haak et al., 2014). Previous normalization reweighting studies involved adapting sequences of around 40 – 60s (Aschner et al., 2018; Yiltiz et al., 2020), but in principle reweighting might occur faster than this, consistent with other types of contrast adaptation.

Multiple suppressive pathways have been identified in the visual system, including between stimuli differing in orientation (Foley, 1994; Heeger, 1992), eye-of-origin (Dougherty et al., 2019; Legge, 1979; Sengpiel & Blakemore, 1994) and spatial position (Cannon & Fullenkamp, 1991; Petrov, 2005). At present there is evidence of normalization reweighting between stimuli with orthogonal orientations (Aschner et al., 2018), and adjacent spatial positions (Yiltiz et al., 2020). We anticipated that interocular suppression should also be subject to reweighting, but that there might be differences in the dynamics across suppressive pathways (Li et al., 2005; e.g. Meese & Baker, 2009; Sengpiel & Vorobyov, 2005). Comparing monocular and dichoptic suppression permits any contribution of early pre-cortical factors to be isolated. This is because interocular suppression is generally thought to impact in primary visual cortex (though see Dougherty et al., 2019), and bypasses any retinal and subcortical stages of processing that contribute to monocular suppression (Li et al., 2005).

Atypical sensory experience, including hypersensitivity to loud sounds, bright lights and strong odours or flavours, is widely reported by individuals on the autism spectrum (MacLennan et al., 2022; Simmons et al., 2009), but the causal mechanisms remain unclear. Fundamental measures of sensitivity including visual acuity (Tavassoli et al., 2011), contrast sensitivity (Koh et al., 2010), and audiometric performance (Rosenhall et al., 1999) are not consistently different from neurotypical controls. Theoretical accounts of sensory differences in autism have proposed that the balance of inhibition and excitation may be disrupted (Rosenberg et al., 2015; Rubenstein & Merzenich, 2003), yet the evidence is currently inconclusive (Sandhu et al., 2020; e.g. Schallmo et al., 2020; Van de Cruys et al., 2018). Our recent work identified an autism-related difference using steady-state EEG (Vilidaite et al., 2018), in which nonlinear (second harmonic) responses were weaker, implicating atypical suppression in autism.

Here we perform a time-course analysis of a previously published data set, and report two novel pre-registered experiments using EEG and MEG. Our data show that suppression increases substantially during the first 2-5 seconds following stimulus onset, for both monocular and dichoptic masks. Source localisation of MEG data indicate that the reweighting is present as early as primary visual cortex (V1). We also hypothesised that normalization reweighting might differ as a function of autistic traits, but did not find convincing support for this hypothesis.

## 3 Materials and Methods

### 3.1 Participants

Experiment 1 was completed by 100 adult participants (32 male, 68 female; mean age 21.9) in early 2015, and first reported by Vilidaite et al. (2018). Here we reanalysed the dataset, and report the results of masking conditions not previously published. Experiment 2 was completed by 100 adult participants (23 male, 74 female, 3 other/not stated; mean age 22.1) in early 2022. Experiment 3 was completed by 10 adults (2 male, 8 female) with a clinical diagnosis of autism, and 10 control participants who were closely matched for age (means of 21.8 and 22, *t* = 0.18, *df* = 18, *p* = 0.86) and exactly matched for gender. Procedures in Experiments 1 and 2 were approved by the ethics committee of the Department of Psychology at the University of York. Procedures for Experiment 3 were approved by the ethics committee of the York Neuroimaging Centre. All participants provided written informed consent, and procedures were consistent with the Declaration of Helsinki.

### 3.2 Apparatus and stimuli

In Experiments 1 and 2, stimuli were presented using a ViewPixx 3D LCD display device (VPixx Technologies, Canada) with a resolution of 1920 × 1080 pixels and a refresh rate of 120Hz. The display was gamma corrected using a Minolta LS110 photometer. In Experiment 2, participants wore active stereo shutter glasses (NVidia 3D Vision 2) that were synchronised with the display using an infra-red signal. EEG data were collected using a 64-channel Waveguard cap, and were amplified and digitised at 1000Hz using an ANT Neuroscan system. Electrode impedance was maintained below 5kΩ, and referenced to a whole-head average.

In Experiment 3, stimuli were presented using a ProPixx DLP projector (VPixx Technologies) running at 120Hz. Stereo presentation was enabled using a circular polariser that was synchronised with the projector refresh, and participants wore passive polarised glasses during the experiment. DLP projectors are perfectly linear, so gamma correction was not required. Data were acquired using a refurbished 248-channel 4D Neuroimaging Magnes 3600 MEG scanner, recording at 1001Hz. Participant head shape was digitised using a Polhemus Fastrak device, and head position was recorded at the start and end of each block by passing current through 5 position coils placed at fiducial points on the head. We also obtained structural MRI scans using a 3 Tesla Siemens Magnetom Prisma scanner to aid in source localisation. Two participants were not available for MRI scans, so we used the MNI ICBM152 template brain (Fonov et al., 2011) for these individuals.

Stimuli were patches of sine wave grating with a diameter of 2 degrees, flickering sinusoidally (on/off flicker) at either 5Hz or 7Hz. In Experiment 1 the gratings had a spatial frequency of 0.5c/deg, and in Experiments 2 & 3 this was increased to 2c/deg. A symmetrical array of 36 individual patches tiled the display. In Experiment 1 the patch orientation was randomly selected on each trial, and all patches had the same orientation. In Experiments 2 & 3 each patch had a random orientation, which was intended to prevent any sequential effects between trials with similar orientations. The central patch was omitted and replaced by a fixation marker constructed from randomly overlaid squares. During each experiment, the fixation marker could be resampled on each trial with a probability of 0.5. Participants were instructed to monitor the fixation marker and count the number of times it changed throughout the experiment. This was intended to maintain attention towards the display and keep participants occupied.

Participants also completed either the short AQ (Hoekstra et al., 2011) in Experiment 1, or the full AQ (Baron-Cohen et al., 2001) in Experiments 2 and 3. For comparison across experiments, we rescaled the short AQ to the same range as the full AQ (0-50). In Experiments 2 and 3, the sensory perception quotient (SPQ) questionnaire (Tavassoli et al., 2014) was also completed.

### 3.3 Experimental design and statistical analysis

In Experiment 1, target stimuli flickering at 7Hz were presented at a range of contrasts (1 - 64%). In half of the conditions a superimposed orthogonal mask of 32% contrast was presented simultaneously, flickering at 5Hz. Stimuli were displayed for trials of 11 seconds, with a 3 second inter-trial interval. The experiment consisted of 4 blocks of trials, each lasting around 10 minutes, and resulting in 8 repetitions of each condition. Participants viewed the display from 57cm, were comfortably seated in an upright position, and were able to rest between blocks. Low latency 8-bit digital triggers transmitted the trial onset and condition information directly to the EEG amplifier.

The procedure for Experiment 2 was very similar, except that participants also wore stereo shutter glasses during the experiment. There were four conditions: (i) monocular presentation of a 5Hz stimulus of 48% contrast, (ii) monocular presentation of a 7Hz stimulus of 48% contrast, (iii) monocular presentation of both stimuli superimposed at right angles, and (iv) dichoptic presentation of both stimuli at right angles (i.e. one stimulus to the left eye, one to the right eye). Eye of presentation was pseudo-randomised to ensure equal numbers of left-eye and right-eye presentations. The trial duration was 6 seconds, with a 3 second inter-trial interval. Participants completed 3 blocks, each lasting around 10 minutes, resulting in a total of 48 repetitions of each condition. Experiment 3 was identical, except that the projector screen was viewed from a distance of 85cm.

EEG data from Experiments 1 and 2 were first imported into Matlab using components of the EEGlab toolbox (Delorme & Makeig, 2004), and converted into a compressed ASCII format. Primary data analysis was then conducted using a bespoke *R* script. In brief, we epoched each trial and extracted the average timecourse across four occipital electrodes (*Oz, POz, O1* and *O2*), and then calculated the Fourier transform of this average waveform. We excluded trials for which the Mahalanobis distance of the complex Fourier components exceeded 3 (for details see Baker, 2021). This resulted in 0.25% of trials being excluded for Experiment 1, and 4.51% of trials for Experiment 2. Next we averaged the waveforms across all remaining trials, and calculated the Fourier transform in a 1-second sliding window to generate timecourses for each participant. We divided the timecourse for the target-only condition by the timecourse for the target + mask condition to produce a suppression ratio. These were then converted to logarithmic (dB) units for averaging, calculation of standard errors, and statistical comparisons. For display purposes we smoothed the timecourses using a cubic spline function, however all statistical comparisons used the unsmoothed data.

Following the suggestion of a reviewer, we conducted an alternative fixed-phase analysis, where the signal in each 1 second epoch was multiplied by a sine wave of appropriate frequency and phase instead of taking the Fourier transform. The results were similar to our main analysis, and can be viewed in the Figures subdirectory of the project code repository (https://github.com/bakerdh/normreweight/tree/main/Figures - see files with the suffix ‘FP’). We also conducted simulations (also available in the code repository) to confirm that our analysis methods were not distorting the estimates of the suppression timecourse. In brief, although the 1 second sliding time window does blur the signals in time, these effects are largely negated by calculating the suppression ratio because the blur cancels out across the numerator and denominator. Overall these simulations give us confidence in the accuracy of our estimates of suppression dynamics.

For Experiment 3, we performed source localisation using a linearly constrained minimum variance (LCMV) beamformer algorithm, implemented in Brainstorm (Tadel et al., 2011). Structural MRI scans were processed using Freesurfer (Dale et al., 1999) to generate a 3D mesh of the head and brain, and we calculated source weights for each block with reference to a 5-minute empty room recording, usually recorded on the day of the experiment. The matrix of source weights for each block was used in a custom Matlab script to extract signals from V1, identified using the probabilistic maps of Wang et al. (2015). These signals were then imported into R for the main analysis, which was consistent with the EEG analysis described above. The outlier rejection procedure excluded 2.47% of trials for Experiment 3.

To make comparisons between groups of participants across time, we used a non-parametric cluster correction technique (Maris & Oostenveld, 2007) based on t-tests. Clusters were identified as temporally adjacent observations that were all statistically significant, and a summed t-value was calculated for each cluster. A null distribution was then generated by randomising group membership and recalculating the summed t-value for the largest cluster, and repeating this procedure 1000 times. Clusters were considered significant if they fell outside of the 95% confidence limits of the null distribution. We adapted this approach to test for significantly increasing suppression by conducting one-way t-tests between time points separated by 1000ms, and repeating the cluster correction procedure as described above.

### 3.4 Preregistration, data and code accessibility

Following a preliminary analysis of the data from Experiment 1, we preregistered our hypotheses and analysis plan for Experiments 2 and 3 on the Open Science Framework website. The preregistration document, along with raw and processed data, and analysis scripts, are publicly available at the project repository: https://osf.io/ab3yv/

## 4 Results

We began by reanalysing data from a steady-state visually evoked potential (SSVEP) experiment reported by Vilidaite et al. (2018). Participants viewed arrays of flickering gratings of varying contrasts. In some conditions a single grating orientation was present flickering at 7Hz (the target), whereas in other conditions a high contrast ‘mask’ was added at right angles to the target gratings, and flickering at 5Hz. The left panel of Figure 1a shows contrast response functions with and without the mask - the presence of the mask reduces the 7Hz response to the target (blue squares are below the black circles; significant main effect of mask contrast, F(1,99) = 26.52, *p* < 0.001). Similarly, the right panel of Figure 1a shows that the 5Hz response to the mask was itself suppressed by the presence of high contrast targets (main effect of target contrast on the mask response, F(2.92,288.63) = 46.77, *p* < 0.001; note that the data from the masking conditions were not reported by Vilidaite et al. (2018)). At both frequencies, responses were localised to the occipital pole (see insets).

**Figure 1:**
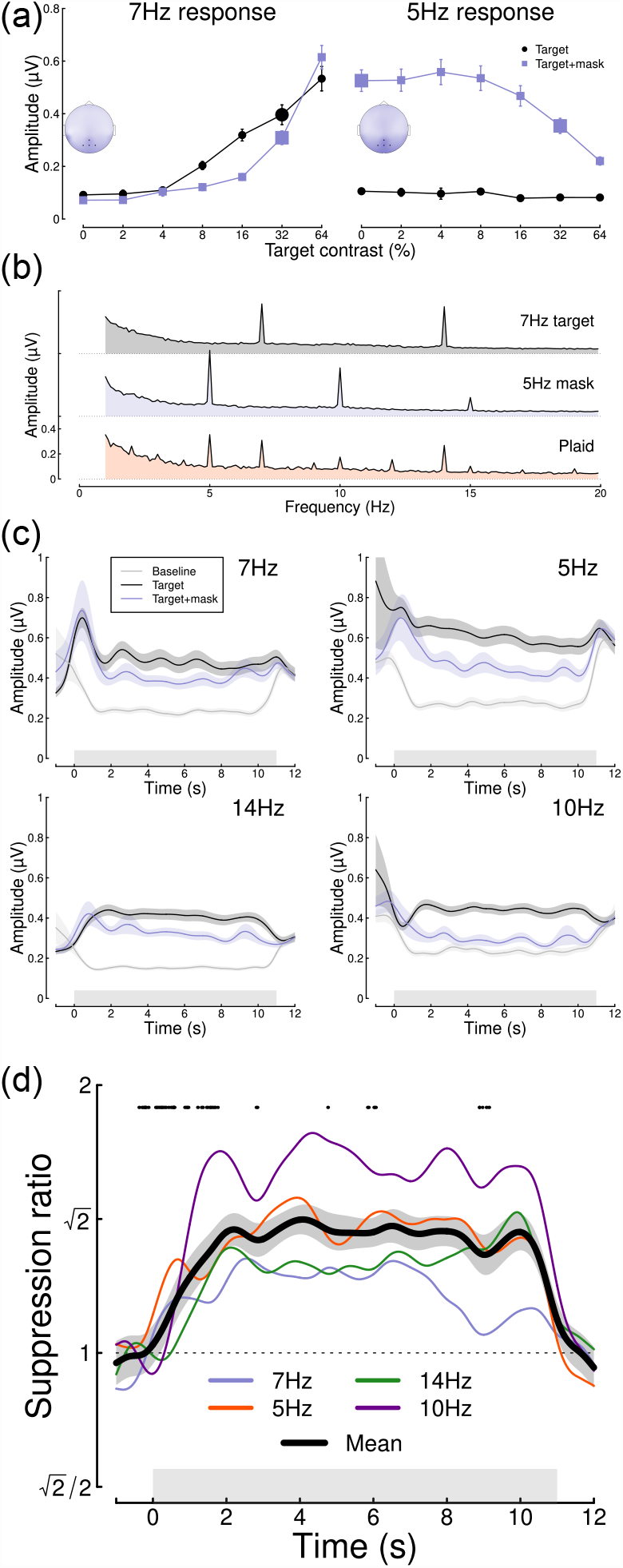
Summary of pilot analysis of data from Vilidiate et al. (2018). Panel (a) shows contrast response functions at the target frequency (7Hz, left) and the mask frequency (5Hz, right). Insets show the distribution of activity across the scalp, with points marking the electrodes over which signals were averaged (Oz, POz, O1 and O2). Panel (b) shows Fourier spectra for the single component stimuli and their combination (plaid). Note the strong second harmonic components at 14Hz and 10Hz. Panel (c) shows timecourses of frequency-locked responses to a single stimulus (black) and the plaid stimulus (blue), compared to a baseline condition (grey) where no stimulus was shown at the target frequency. Panel (d) shows the timecourse of suppression at each frequency (7Hz, 5Hz, 14Hz, 10Hz) and their average (black curve). Points around y = 1.8 indicate a significantly increasing ratio (for the time window centred at each point). Error bars in panel (a) and shaded regions in panels (c,d) indicate ±1SE across N=100 participants, and grey rectangles indicate the timing of stimulus presentation. The larger symbols in panel (a) indicate the conditions used for subsequent analyses.

We then performed a timecourse analysis, in which we analysed each 11-second trial using a sliding 1-second time window. The top panel of Figure 1c shows the response at the target frequency (7Hz) to a single stimulus of 32% contrast (black), and the response at 7Hz when the 32% contrast mask is added (blue). For comparison, a baseline timecourse is also shown (grey), which was the target response at 7Hz when a 5Hz mask stimulus was shown (therefore controlling for attention, blinking etc.). Analogous responses are shown at three other frequencies - the mask frequency (5Hz), and the second harmonics of both target and mask frequencies (14Hz, 10Hz), at which strong responses were also found (see spectra in Figure 1b). The reduction in signal strength when a second component is added at a different orientation and frequency illustrates the masking effect. Surprisingly there was sometimes substantial activity before and after the stimulus was presented, as is especially clear in the baseline condition shown by the grey curves in Figure 1c. We think the most likely explanation for this is broadband noise from participant movement during the breaks between trials. Since it is approximately equal across conditions it appears to cancel out in the suppression ratios (Figure 1d).

Taking the ratio of the two timecourses (the target only timecourse and the target timecourse when a mask was present) to calculate a masking index reveals that for 7Hz targets masking increases steeply during the first two seconds of stimulus presentation, and then plateaus for several seconds (blue trace in Figure 1d). A similar pattern is observed for the 5Hz mask (red trace in Figure 1d), as well as at the second harmonics, with some variability in the timecourse across frequencies; for example, at 5Hz suppression peaks at around 4 seconds. The black trace shows the average masking ratio across all four frequencies, which rises steeply for just over two seconds and then stays approximately constant until stimulus offset. We conducted cluster-corrected t-tests between ratios separated by 1000ms, testing for an increase in suppression ratio across time (i.e. a one-sided test). Points at y = 1.8 in Figure 1d indicate time points where the ratio is significantly increasing (i.e. there is significantly more suppression 500ms after the time point than there was 500ms before it), and occur up until 2.27 seconds after stimulus presentation. We also calculated an overall effect size by comparing the amount of suppression during the first 1000ms following stimulus onset with that between 2000 and 3000ms, averaged across all temporal frequencies. This effect size (Cohen’s *d* = 0.49) indicated a medium-sized effect.

Our initial reanalysis was promising, however the data were noisy despite the large sample size (of N=100), because each participant contributed only 8 trials (88 seconds) to each condition. We therefore preregistered two new experiments (see https://osf.io/4qudc) to investigate these effects in greater detail. These had a similar overall design to the Vilidaite et al. (2018) study, with some small changes intended to optimise the study (see Methods). The key differences were that we used shorter trials (because there were few changes in the latter part of the trials shown in Figure 1d), and also focussed all trials into a smaller number of conditions, such that each participant contributed 48 repetitions (288 seconds of data) to each of 4 conditions.

Figure 2 summarises the results of our EEG experiment testing a further 100 adult participants. Averaged EEG waveforms showed a strong oscillatory component at each of the two stimulus flicker frequencies (Figure 2a), which slightly lagged the driving signal. Signals were well-isolated in the Fourier domain (Figure 2b), and localised to occipital electrodes. Responses at 7Hz were weaker in the two masking conditions, showing significant changes in response amplitude for both the monocular (*t* = 7.56, *df* = 87, *p* < 0.001) and dichoptic (*t* = 11.35, *df* = 87, *p* < 0.001) masks. Dichoptic masking was significantly stronger than monocular masking (*t* = 7.96, *df* = 87, *p* < 0.001), and a similar pattern was evident at 5Hz (note that for this experiment, the terms ‘target’ and ‘mask’ are arbitrary, as each component was presented at a single contrast).

**Figure 2:**
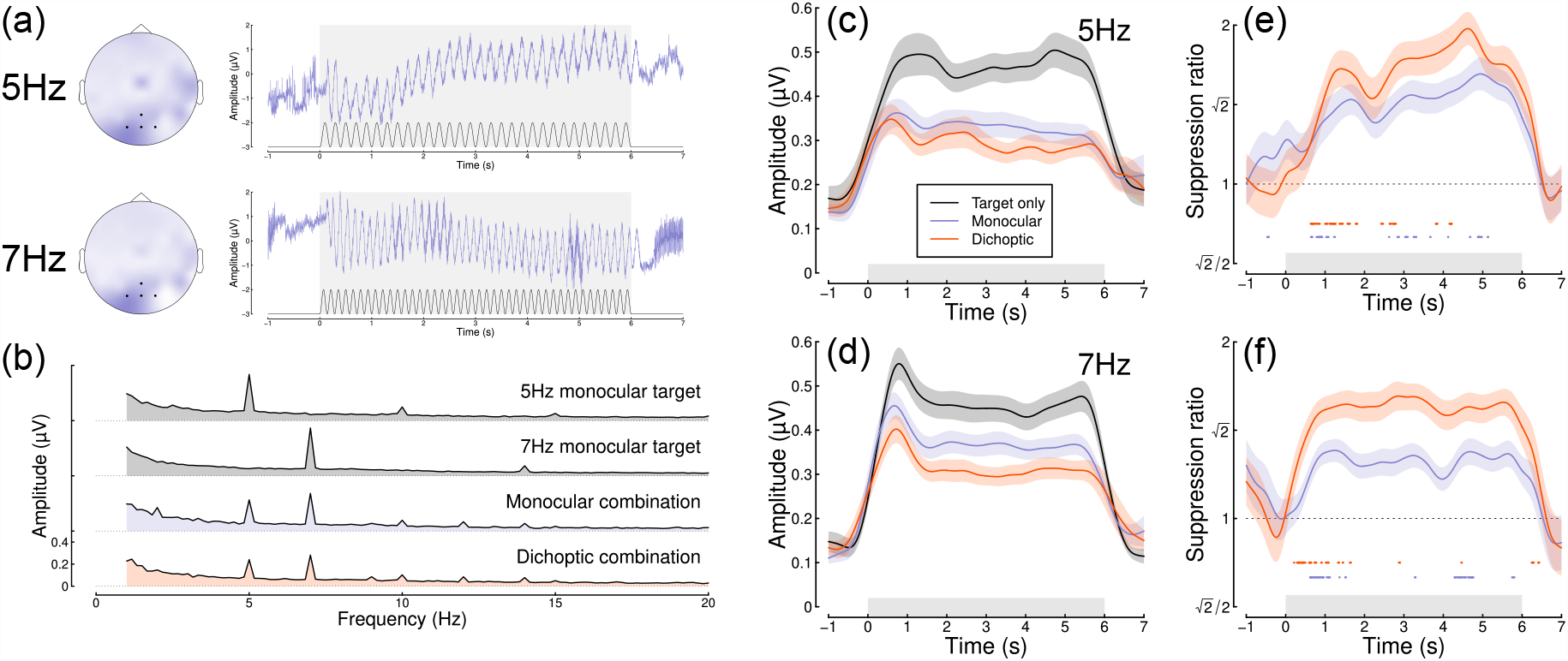
Summary of EEG results for N=100 adult participants. Panel (a) shows scalp topographies and averaged waveforms for 5Hz (top) and 7Hz (bottom) stimuli. The black sine wave trace in each panel illustrates the driving contrast modulation, and black points on the scalp topographies indicate electrodes Oz, O1, O2 and POz. Panel (b) shows the Fourier amplitude spectrum for each condition, with clear peaks at 5Hz and 7Hz. Panels (c,d) show timecourses at each frequency for the target-only condition (black), and the monocular (blue) and dichoptic (red) masking conditions. Panels (e,f) show suppression ratios as a function of time for each mask type, with points around y = 0.8 indicating a significantly increasing ratio. Shaded regions in panels (c-f) span ±1SE across participants, and light grey rectangles indicate the period of stimulus presentation.

The timecourse at both flicker frequencies showed an initial onset transient, and was then relatively stable for the 6 seconds of stimulus presentation (Figure 2c,d). The ratio of target only to target + mask conditions increased over time (Figure 2e,f) for both mask types. At 5Hz the increase in masking continued for as long as 5 seconds of stimulus presentation in the monocular condition (Figure 2e; points at y = 0.8 indicate significantly increasing suppression, which continue until 5.1s (mon) or 4.2s (dich)), whereas at 7Hz the increase occurred primarily during the first 1.5 seconds after onset (Figure 2f; substantial clusters up to 1.5s (mon) and 1.7s (dich)). These differences across frequency are consistent with the pilot data (see Figure 1d). Both monocular and dichoptic masks produced similar timecourses of suppression. We calculated an overall effect size comparing suppression in the first 1000ms after stimulus onset to the time window from 3000-4000ms, pooling over frequency and mask type. This had a value of *d* = 0.33. Overall, this second study confirmed that normalization increases during the first few seconds of a steady-state trial, and extends this finding to dichoptic mask arrangements.

Next we repeated the experiment on 20 participants using a 248-channel whole-head cryogenic MEG system. Half of the participants had a diagnosis of autism, and the remainder were age- and gender-matched controls. Source localisation using a linearly constrained minimum variance (LCMV) beamformer algorithm (Van Veen et al., 1997) showed strong localisation of steady-state signals at the occipital pole (see Figure 3a), and in the Fourier domain (Figure 3b). Responses from the most responsive V1 vertex showed a similar timecourse to those of the EEG experiments at both frequencies (Figure 3c,d), and showed increasing suppression during the first few seconds of stimulus presentation (Figure 3e,f). The normalization reweighting effect was again clearest at 5Hz, especially for the dichoptic condition (red curve in Figure 3e), which increased until 2.5s. This confirms that the reweighting effects can occur as early as primary visual cortex, consistent with findings from neurophysiology (Aschner et al., 2018). However the data are more variable than for our EEG experiments, and had fewer significant clusters, perhaps owing to a power reduction caused by the smaller sample size for this dataset and/or greater heterogeneity across frequency. When pooling effects over frequency and condition, the overall effect size (*d* = 0.03) was near zero.

**Figure 3:**
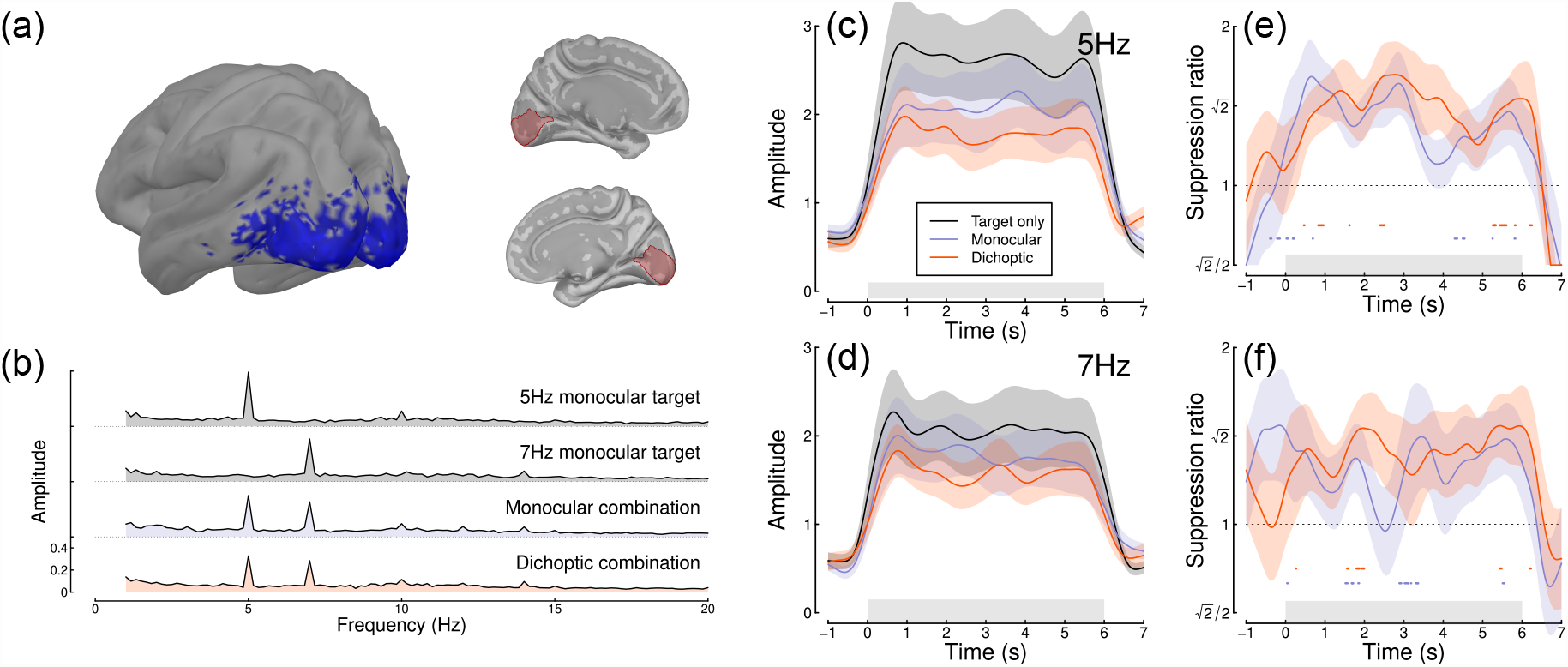
Summary of MEG results for N=20 adults. Panel (a) shows average SSVEP response in source space, thresholded at SNR=2 (blue, left), and locations of the V1 ROI on the medial surface of both hemispheres (right, red). Panel (b) shows the Fourier spectra for the four experimental conditions, from the most responsive vertex in V1. Panels (c,d) show timecourses at 5Hz and 7Hz, and panels (e,f) show suppression ratios for the monocular and dichoptic conditions at each frequency, with points around y = 0.8 indicating a significantly increasing ratio. Shaded regions in panels (c-f) indicate ±1SE across participants, and light grey rectangles indicate the period of stimulus presentation.

Intermodulation responses, at sums and differences of different stimulation frequencies, are another marker of nonlinear interaction (Cunningham et al., 2017; Regan & Regan, 1988; Tsai et al., 2012). We also calculated the timecourse of the sum intermodulation terms (at 12Hz) in our data sets (the difference terms at 2Hz were negligible). Figure 4 shows that for both EEG experiments, the intermodulation term increases during the first 1 second of stimulus presentation and then remains approximately constant. The intermodulation response in the MEG data was less clear, consistent with the spectra shown in Figure 3b. It seems unlikely that intermodulation terms are useful for monitoring the timecourse of normalization reweighting, and indeed they may derive from a nonlinear process other than suppression, such as exponentiation and signal combination (Regan & Regan, 1988). Previous work has identified situations in which suppression is constant, but the intermodulation term changes substantially between conditions depending on the extent of signal pooling (Cunningham et al., 2017).

**Figure 4:**
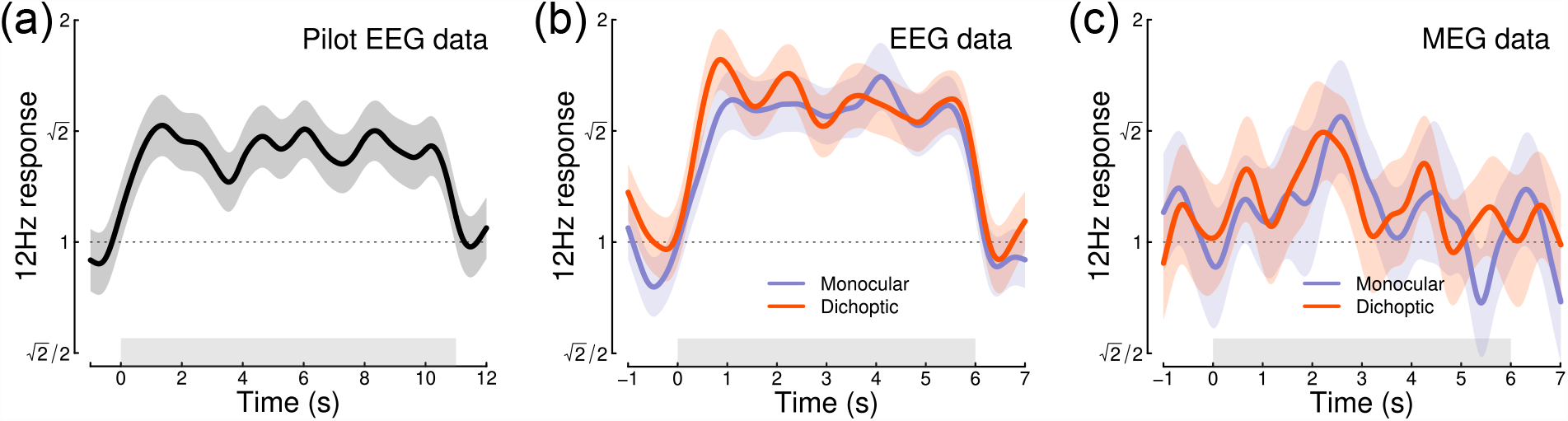
Timecourse of the sum intermodulation term at 12Hz across three experiments. In both EEG experiments, the intermodulation response increases during the first 1 second of stimulus presentation. Responses are calculated as a proportional increase relative to the target-only conditions (where the intermodulation response is absent), for direct comparison with the suppression ratios in Figures 1-3. Shaded regions indicate ±1SE.

To investigate whether normalization reweighting effects differ with respect to autistic traits, we then split each dataset (averaged across temporal frequency) using median AQ score (for the EEG experiments) or according to diagnostic group (autism vs controls) for the MEG data. Figure 5a-c shows distributions of AQ scores for each experiment, and indicates for the pilot and EEG data which participants were in the high (purple) and low (green) AQ groups. The median AQ scores were 14 for the pilot data, and 18 for the EEG data. In the MEG experiment, AQ scores for the autism group (mean 36.1) and the control group (mean 16.7) were significantly different (*t* = 6.00, *df* = 14.2, *p* < 0.001), with minimal overlap (one participant with an autism diagnosis had an AQ score marginally lower than the highest AQ scores from the control group). These distributions are consistent with previous results for AQ (Baron-Cohen et al., 2001).

**Figure 5:**
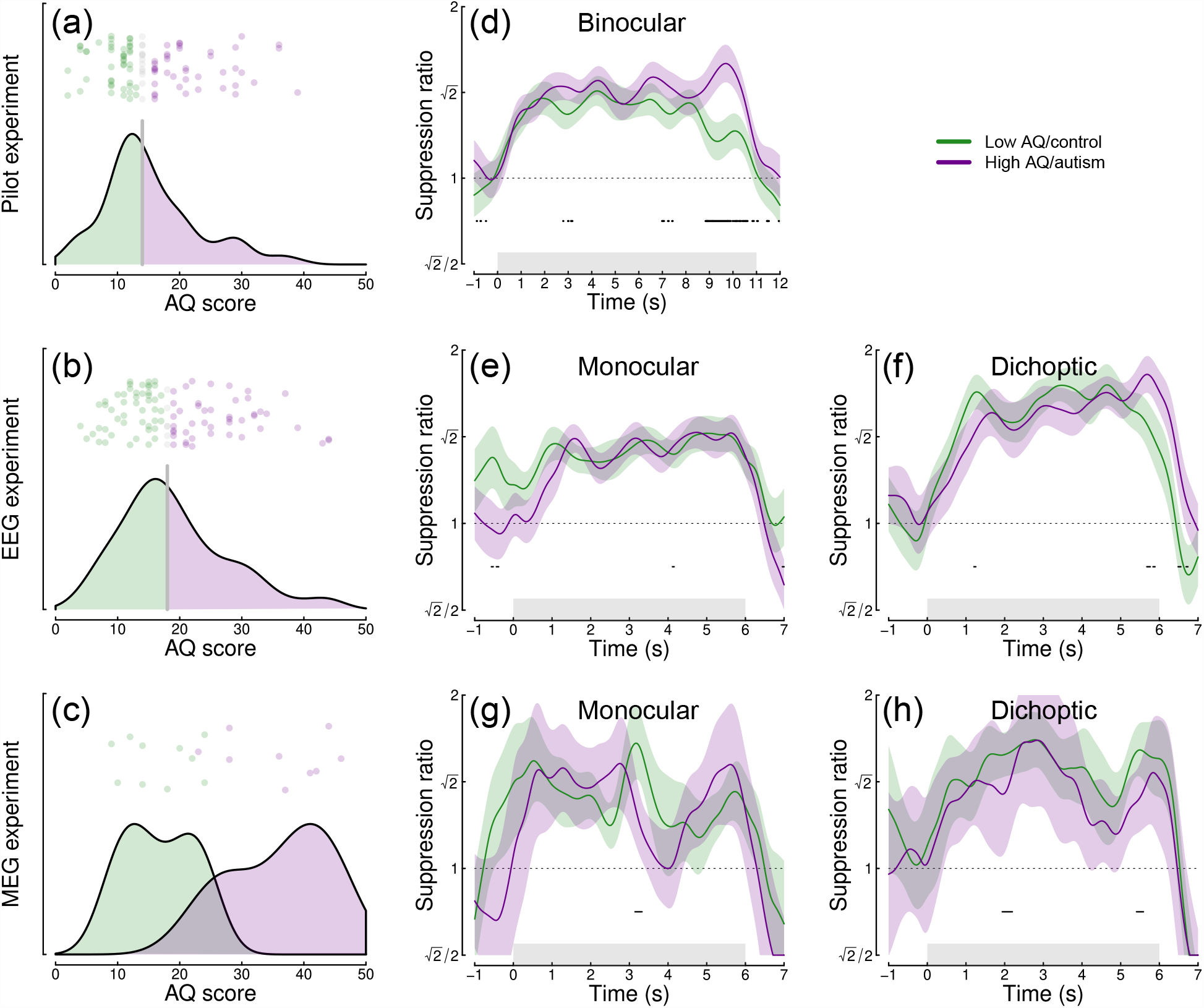
Analysis of the effect of autistic traits on normalization reweighting. Panels (a-c) show distributions of AQ scores across the three data sets. Panels (d-h) show timecourses of suppression averaged across stimulation frequency, and split by AQ score (d-f) or autism status (g, h). Panels (d,e,g) are for binocular or monocular presentation, and panels (f,h) are for dichoptic presentation. Shaded regions in panels (d-h) indicate ±1SE across participants, and black points at y = 0.8 indicate significant differences between groups.

We compared the timecourse of suppression between groups using a nonparametric cluster correction approach (Maris & Oostenveld, 2007) to control the type I error rate. Significant clusters are indicated at y = 0.8 in panels d-h of Figure 5. Despite some occasionally significant clusters, there is no clear or consistent difference between groups across our three data sets. In particular, none of the significant clusters occur during the first few seconds of stimulus onset, when reweighting takes place. We also compared suppression ratios calculated on Fourier components for the full trial, and found no significant effects of autism on suppression strength. For Experiment 1 we assessed the first and second harmonics separately, but also found no AQ-related differences. We therefore conclude that autism/AQ score is not associated with normalization reweighting, or the strength of suppression more generally.

## 5 Discussion

We found evidence of dynamic normalization reweighting across three separate datasets. Suppression increased significantly during the first 2-5 seconds of stimulus presentation, though with some variation across temporal frequency. Relative to the first 1 second of stimulus presentation, the increase in suppression after 3 seconds constituted an effect size of *d* = 0.49 for the pilot data, *d* = 0.33 for our new EEG experiment, and *d* = 0.03 for our MEG experiment (pooled across temporal frequency and monocular and dichoptic conditions). Reweighting had a similar timecourse for monocular and dichoptic stimulus presentation, and was apparent as early as V1. We did not find compelling differences associated with autism, or high vs low autistic traits. In the remainder of this section we will discuss possible explanations for temporal frequency differences, evidence for inhibitory differences in autism, and more general implications of dynamic normalization reweighting.

One important question is whether the dynamic increase in suppression can be explained by the stimulus onset transient. This is a possibility that cannot be ruled out for some of our data. For example, the steep increase in suppression in Figure 2f has a similar timecourse to the onset transient in Figure 2d. However, there are also counterexamples where suppression continues to increase well beyond the first 1 second of stimulus presentation (e.g. Figure 2e). It is currently unclear why there appear to be such substantial differences between temporal frequency conditions, especially with such similar frequencies (5 and 7Hz). However the differences are relatively consistent across experiments. For example, 5Hz flicker produces a more gradual increase in suppression across all three data sets, compared with 7Hz flicker. These differences may be a consequence of visual channels with different temporal tuning interacting with the stimulation frequency, as well as any nonlinearities that govern suppression. Or there could be an asymmetry, whereby the relative temporal frequency between the two stimulus components affects the character of suppression (Liza & Ray, 2022). We hope to be able to model these effects in the future, for example by using dynamic models of early vision that incorporate time-lagged gain control (e.g. Zhou et al., 2019).

We did not observe clear differences in the timecourse between monocular and dichoptic suppression. This is important, because the dichoptic arrangement bypasses early stages of processing before the cortex (e.g. the retina and lateral geniculate nucleus). It suggests that the dynamic increases in suppression occur in the cortex, consistent with our MEG data that find evidence of reweighting in V1 (see Fig 3), and with previous neurophysiological work (Aschner et al., 2018). It is currently unclear whether these effects originate in V1, or might involve feedback from higher areas. The similarity between monocular and dichoptic effects also differs from work on adaptation to individual mask components. In both physiological (Li et al., 2005; Sengpiel & Vorobyov, 2005) and psychophysical (Baker et al., 2007) paradigms, adapting to a dichoptic mask reduces its potency, whereas adapting to a monocular mask has little or no effect. Normalization reweighting offers an explanation for why monocular masks presented in isolation do not adapt: if suppressive weights are determined by co-occurrence of stimuli, presentation of an isolated mask will have little effect. However this cannot explain the dichoptic adaptation effects without invoking additional binocular processes, such as competition between summing and differencing channels (e.g. May et al., 2012).

The relationship between normalization reweighting and other forms of visual plasticity and adaptation is currently unclear. One phenomenon that might be closely related to our dichoptic effect is the change in interocular suppression that occurs when one eye is patched for a period of time (Lunghi et al., 2011). In the patching paradigm, the inputs to the two eyes are uncorrelated while one eye is patched, which the normalization reweighting model predicts should reduce suppression between the eyes. Most studies using patching have focussed on the resulting imbalance between the patched and non-patched eye, in which the patched eye contributes more to binocular single vision than the non-patched eye. In principle this could be due to increased suppression of the non-patched eye (inconsistent with normalization reweighting), or reduced suppression of the patched eye (consistent with normalization reweighting). It is difficult to distinguish these possibilities using paradigms that assess the balance between the two eyes, such as the binocular rivalry paradigm from the original Lunghi et al. (2011) study. However subsequent work has shown that patching increases the patched eye’s response (Zhou et al., 2015), and reduces both dichoptic masking (Baldwin & Hess, 2018) and levels of the inhibitory neurotransmitter GABA (Lunghi et al., 2015). All of these findings are consistent with a reweighting account.

Autism is composed of a set of heterogenous symptoms and characteristics, and normalization reweighting may have a more specific relationship to some aspects of autism, rather than autism per se. For this reason, we also examined relationships with the sensory perception quotient (SPQ) to examine whether sensory experiences specifically were related to normalization reweighting. SPQ scores showed significant negative correlation with AQ for the data sets from Experiments 2 and 3 (EEG data, *r* = -0.35, *p* < 0.001; MEG data, *r* = -0.57, *p* = 0.011) with effect sizes comparable to those reported previously (Tavassoli et al., 2014). We also conducted an exploratory analysis of the EEG data from Experiment 2, splitting participants by SPQ instead of AQ. However this analysis did not reveal any convincing differences in normalization reweighting either. Our preregistration also proposed to replicate our earlier finding of a reduced second harmonic response in participants with autism/high AQ scores. However the changes to the experimental design greatly reduced the second harmonic response in both experiments, such that it could not be observed reliably (see Figures 2b and 3b). We were therefore not confident in conducting this analysis. We suspect that the increase in spatial frequency from 0.5 c/deg in the Vilidaite et al. (2018) study to 2 c/deg here is most likely responsible for the dramatically reduced second harmonic response.

The idea that the dynamic balance of inhibition and excitation might be different in autism (Rosenberg et al., 2015; Rubenstein & Merzenich, 2003) has compelling face validity. For example, individuals with autism often report difficulties with changes in their sensory environment, which might be due to gain control processes failing to adapt appropriately. Indeed, there is experimental evidence of reduced adaptation across various domains (Pellicano et al., 2007; Turi et al., 2015), which is predicted by some autism models (Pellicano & Burr, 2012). However this appears not to extend to changes in normalization reweighting, despite the link between reweighting and adaptation (Westrick et al., 2016).

### 5.1 Conclusions

We investigated the timecourse of normalization reweighting across three datasets, with a total of 220 participants. We found clear evidence that suppression increases during the first 2-5 seconds of stimulus presentation, though there were differences across frequency that are currently unexplained. We did not find evidence of autism-related differences in either the magnitude or timecourse of suppression. Our results support an emerging theory that suppression is a dynamic process that allows sensory systems to recalibrate according to their recent history.

## 6 Acknowledgements

Supported by BBSRC grant BB/V007580/1 awarded to DHB and ARW. We are grateful to all of the participants who took part in the experiments reported here.

## References

Aschner, A., Solomon, S. G., Landy, M. S., Heeger, D. J., & Kohn, A. (2018). Temporal contingencies determine whether adaptation strengthens or weakens normalization. J Neurosci, 38 (47), 10129–10142. https://doi.org/10.1523/JNEUROSCI.1131-18.2018

Baker, D. H. (2021). Statistical analysis of periodic data in neuroscience. Neurons, Behavior, Data Analysis, and Theory, 5 (3). https://doi.org/10.51628/001c.27680

Baker, D. H., Meese, T. S., & Summers, R. J. (2007). Psychophysical evidence for two routes to suppression before binocular summation of signals in human vision. Neuroscience, 146 (1), 435–448. https://doi.org/10.1016/j.neuroscience.2007.01.030

Baldwin, A. S., & Hess, R. F. (2018). The mechanism of short-term monocular deprivation is not simple: Separate effects on parallel and cross-oriented dichoptic masking. Sci Rep, 8 (1), 6191. https://doi.org/10.1038/s41598-018-24584-9

Baron-Cohen, S., Wheelwright, S., Skinner, R., Martin, J., & Clubley, E. (2001). The autism-spectrum quotient (AQ): Evidence from asperger syndrome/high-functioning autism, males and females, scientists and mathematicians. J Autism Dev Disord, 31 (1), 5–17. https://doi.org/10.1023/a:1005653411471

Blakemore, C., & Campbell, F. W. (1969). On the existence of neurones in the human visual system selectively sensitive to the orientation and size of retinal images. J Physiol, 203 (1), 237–260. https://doi.org/10.1113/jphysiol.1969.sp008862

Cannon, M. W., & Fullenkamp, S. C. (1991). Spatial interactions in apparent contrast: Inhibitory effects among grating patterns of different spatial frequencies, spatial positions and orientations. Vision Res, 31 (11), 1985–1998. https://doi.org/10.1016/0042-6989(91)90193-9

Carandini, M., & Heeger, D. J. (2011). Normalization as a canonical neural computation. Nature Reviews Neuroscience, 13 (1), 51–62. https://doi.org/10.1038/nrn3136

Cunningham, D. G. M., Baker, D. H., & Peirce, J. W. (2017). Measuring nonlinear signal combination using EEG. J Vis, 17 (5), 10. https://doi.org/10.1167/17.5.10

Dale, A. M., Fischl, B., & Sereno, M. I. (1999). Cortical surface-based analysis. I. Segmentation and surface reconstruction. Neuroimage, 9 (2), 179–194. https://doi.org/10.1006/nimg.1998.0395

Delorme, A., & Makeig, S. (2004). EEGLAB: An open source toolbox for analysis of single-trial EEG dynamics including independent component analysis. J Neurosci Methods, 134 (1), 9–21. https://doi.org/10.1016/j.jneumeth.2003.10.009

Dougherty, K., Schmid, M. C., & Maier, A. (2019). Binocular response modulation in the lateral geniculate nucleus. J Comp Neurol, 527 (3), 522–534. https://doi.org/10.1002/cne.24417

Foley, J. M. (1994). Human luminance pattern-vision mechanisms: Masking experiments require a new model. J Opt Soc Am A Opt Image Sci Vis, 11 (6), 1710–1719. https://doi.org/10.1364/josaa.11.001710

Foley, J. M., & Chen, C. C. (1997). Analysis of the effect of pattern adaptation on pattern pedestal effects: A two-process model. Vision Res, 37 (19), 2779–2788. https://doi.org/10.1016/s0042-6989(97)00081-3

Fonov, V., Evans, A. C., Botteron, K., Almli, C. R., McKinstry, R. C., Collins, D. L., & Brain Development Cooperative Group. (2011). Unbiased average age-appropriate atlases for pediatric studies. Neuroimage, 54 (1), 313–327. https://doi.org/10.1016/j.neuroimage.2010.07.033

Freeman, T. C. B., Durand, S., Kiper, D. C., & Carandini, M. (2002). Suppression without inhibition in visual cortex. Neuron, 35 (4), 759–771. https://doi.org/10.1016/s0896-6273(02)00819-x

Gibson, J. J., & Radner, M. (1937). Adaptation, after-effect and contrast in the perception of tilted lines. I. Quantitative studies. Journal of Experimental Psychology, 20 (5), 453–467. https://doi.org/10.1037/h0059826

Greenlee, M. W., Georgeson, M. A., Magnussen, S., & Harris, J. P. (1991). The time course of adaptation to spatial contrast. Vision Res, 31 (2), 223–236. https://doi.org/10.1016/0042-6989(91)90113-j

Haak, K. V., Fast, E., Bao, M., Lee, M., & Engel, S. A. (2014). Four days of visual contrast deprivation reveals limits of neuronal adaptation. Curr Biol, 24 (21), 2575–2579. https://doi.org/10.1016/j.cub.2014.09.027

Heeger, D. J. (1992). Normalization of cell responses in cat striate cortex. Vis Neurosci, 9 (2), 181–197. https://doi.org/10.1017/s0952523800009640

Hoekstra, R. A., Vinkhuyzen, A. A. E., Wheelwright, S., Bartels, M., Boomsma, D. I., Baron-Cohen, S., Posthuma, D., & Sluis, S. van der. (2011). The construction and validation of an abridged version of the autism-spectrum quotient (AQ-short). J Autism Dev Disord, 41 (5), 589–596. https://doi.org/10.1007/s10803-010-1073-0

Koh, H. C., Milne, E., & Dobkins, K. (2010). Spatial contrast sensitivity in adolescents with autism spectrum disorders. J Autism Dev Disord, 40 (8), 978–987. https://doi.org/10.1007/s10803-010-0953-7

Kwon, M., Legge, G. E., Fang, F., Cheong, A. M. Y., & He, S. (2009). Adaptive changes in visual cortex following prolonged contrast reduction. J Vis, 9 (2), 20.1–16. https://doi.org/10.1167/9.2.20

Legge, G. E. (1979). Spatial frequency masking in human vision: Binocular interactions. J Opt Soc Am, 69 (6), 838–847. https://doi.org/10.1364/josa.69.000838

Li, B., Peterson, M. R., Thompson, J. K., Duong, T., & Freeman, R. D. (2005). Cross-orientation suppression: Monoptic and dichoptic mechanisms are different. J Neurophysiol, 94 (2), 1645–1650. https://doi.org/10.1152/jn.00203.2005

Liza, K., & Ray, S. (2022). Local interactions between steady-state visually evoked potentials at nearby flickering frequencies. J Neurosci, 42 (19), 3965–3974. https://doi.org/10.1523/JNEUROSCI.0180-22.2022

Lunghi, C., Burr, D. C., & Morrone, C. (2011). Brief periods of monocular deprivation disrupt ocular balance in human adult visual cortex. Curr Biol, 21 (14), R538–9. https://doi.org/10.1016/j.cub.2011.06.004

Lunghi, C., Emir, U. E., Morrone, M. C., & Bridge, H. (2015). Short-term monocular deprivation alters GABA in the adult human visual cortex. Curr Biol, 25 (11), 1496–1501. https://doi.org/10.1016/j.cub.2015.04.021

MacLennan, K., O’Brien, S., & Tavassoli, T. (2022). In our own words: The complex sensory experiences of autistic adults. J Autism Dev Disord, 52 (7), 3061–3075. https://doi.org/10.1007/s10803-021-05186-3

Maris, E., & Oostenveld, R. (2007). Nonparametric statistical testing of EEG- and MEG-data. J Neurosci Methods, 164 (1), 177–190. https://doi.org/10.1016/j.jneumeth.2007.03.024

Mather, G., Pavan, A., Campana, G., & Casco, C. (2008). The motion aftereffect reloaded. Trends Cogn Sci, 12 (12), 481–487. https://doi.org/10.1016/j.tics.2008.09.002

May, K. A., Zhaoping, L., & Hibbard, P. B. (2012). Perceived direction of motion determined by adaptation to static binocular images. Curr Biol, 22 (1), 28–32. https://doi.org/10.1016/j.cub.2011.11.025

Meese, T. S., & Baker, D. H. (2009). Cross-orientation masking is speed invariant between ocular pathways but speed dependent within them. Journal of Vision, 9 (5), 2. https://doi.org/10.1167/9.5.2

Pellicano, E., & Burr, D. (2012). When the world becomes ‘too real’: A bayesian explanation of autistic perception. Trends Cogn Sci, 16 (10), 504–510. https://doi.org/10.1016/j.tics.2012.08.009

Pellicano, E., Jeffery, L., Burr, D., & Rhodes, G. (2007). Abnormal adaptive face-coding mechanisms in children with autism spectrum disorder. Curr Biol, 17 (17), 1508–1512. https://doi.org/10.1016/j.cub.2007.07.065

Petrov, Y. (2005). Two distinct mechanisms of suppression in human vision. Journal of Neuroscience, 25 (38), 8704–8707. https://doi.org/10.1523/jneurosci.2871-05.2005

Regan, M. P., & Regan, D. (1988). A frequency domain technique for characterizing nonlinearities in biological systems. Journal of Theoretical Biology, 133 (3), 293–317. https://doi.org/https://doi.org/10.1016/S0022-5193(88)80323-0

Rosenberg, A., Patterson, J. S., & Angelaki, D. E. (2015). A computational perspective on autism. Proc Natl Acad Sci U S A, 112 (30), 9158–9165. https://doi.org/10.1073/pnas.1510583112

Rosenhall, U., Nordin, V., Sandström, M., Ahlsén, G., & Gillberg, C. (1999). Autism and hearing loss. J Autism Dev Disord, 29 (5), 349–357. https://doi.org/10.1023/a:1023022709710

Rubenstein, J. L. R., & Merzenich, M. M. (2003). Model of autism: Increased ratio of excitation/inhibition in key neural systems. Genes Brain Behav, 2 (5), 255–267. https://doi.org/10.1034/j.1601-183x.2003.00037.x

Sandhu, T. R., Reese, G., & Lawson, R. P. (2020). Preserved low-level visual gain control in autistic adults. Wellcome Open Research, 4 (208). https://doi.org/10.12688/wellcomeopenres.15615.1

Schallmo, M.-P., Kolodny, T., Kale, A. M., Millin, R., Flevaris, A. V., Edden, R. A. E., Gerdts, J., Bernier, R. A., & Murray, S. O. (2020). Weaker neural suppression in autism. Nat Commun, 11 (1), 2675. https://doi.org/10.1038/s41467-020-16495-z

Sengpiel, F., & Blakemore, C. (1994). Interocular control of neuronal responsiveness in cat visual cortex. Nature, 368 (6474), 847–850. https://doi.org/10.1038/368847a0

Sengpiel, F., & Vorobyov, V. (2005). Intracortical origins of interocular suppression in the visual cortex. J Neurosci, 25 (27), 6394–6400. https://doi.org/10.1523/JNEUROSCI.0862-05.2005

Simmons, D. R., Robertson, A. E., McKay, L. S., Toal, E., McAleer, P., & Pollick, F. E. (2009). Vision in autism spectrum disorders. Vision Research, 49 (22), 2705–2739. https://doi.org/10.1016/j.visres.2009.08.005

Tadel, F., Baillet, S., Mosher, J. C., Pantazis, D., & Leahy, R. M. (2011). Brainstorm: A user-friendly application for MEG/EEG analysis. Comput Intell Neurosci, 2011, 879716. https://doi.org/10.1155/2011/879716

Tavassoli, T., Hoekstra, R. A., & Baron-Cohen, S. (2014). The sensory perception quotient (SPQ): Development and validation of a new sensory questionnaire for adults with and without autism. Mol Autism, 5, 29. https://doi.org/10.1186/2040-2392-5-29

Tavassoli, T., Latham, K., Bach, M., Dakin, S. C., & Baron-Cohen, S. (2011). Psychophysical measures of visual acuity in autism spectrum conditions. Vision Research, 51 (15), 1778–1780. https://doi.org/10.1016/j.visres.2011.06.004

Tsai, J. J., Wade, A. R., & Norcia, A. M. (2012). Dynamics of normalization underlying masking in human visual cortex. J Neurosci, 32 (8), 2783–2789. https://doi.org/10.1523/JNEUROSCI.4485-11.2012

Turi, M., Burr, D. C., Igliozzi, R., Aagten-Murphy, D., Muratori, F., & Pellicano, E. (2015). Children with autism spectrum disorder show reduced adaptation to number. Proc Natl Acad Sci U S A, 112 (25), 7868–7872. https://doi.org/10.1073/pnas.1504099112

Van de Cruys, S., Vanmarcke, S., Steyaert, J., & Wagemans, J. (2018). Intact perceptual bias in autism contradicts the decreased normalization model. Scientific Reports, 8 (1). https://doi.org/10.1038/s41598-018-31042-z

Van Veen, B. D., Drongelen, W. van Yuchtman, M., & Suzuki, A. (1997). Localization of brain electrical activity via linearly constrained minimum variance spatial filtering. IEEE Trans Biomed Eng, 44 (9), 867–880. https://doi.org/10.1109/10.623056

Vilidaite, G., Norcia, A. M., West, R. J. H., Elliott, C. J. H., Pei, F., Wade, A. R., & Baker, D. H. (2018). Autism sensory dysfunction in an evolutionarily conserved system. Proc Biol Sci, 285 (1893), 20182255. https://doi.org/10.1098/rspb.2018.2255

Wang, L., Mruczek, R. E. B., Arcaro, M. J., & Kastner, S. (2015). Probabilistic maps of visual topography in human cortex. Cereb Cortex, 25 (10), 3911–3931. https://doi.org/10.1093/cercor/bhu277

Webster, M. A. (2015). Visual adaptation. Annu Rev Vis Sci, 1, 547–567. https://doi.org/10.1146/annurev-vision-082114-035509

Westrick, Z. M., Heeger, D. J., & Landy, M. S. (2016). Pattern adaptation and normalization reweighting. Journal of Neuroscience, 36 (38), 9805–9816. https://doi.org/10.1523/jneurosci.1067-16.2016

Yiltiz, H., Heeger, D. J., & Landy, M. S. (2020). Contingent adaptation in masking and surround suppression. Vision Res, 166, 72–80. https://doi.org/10.1016/j.visres.2019.11.004

Zhou, J., Baker, D. H., Simard, M., Saint-Amour, D., & Hess, R. F. (2015). Short-term monocular patching boosts the patched eye’s response in visual cortex. Restor Neurol Neurosci, 33 (3), 381–387. https://doi.org/10.3233/RNN-140472

Zhou, J., Benson, N. C., Kay, K., & Winawer, J. (2019). Predicting neuronal dynamics with a delayed gain control model. PLoS Comput Biol, 15 (11), e1007484. https://doi.org/10.1371/journal.pcbi.1007484

